# A pilot study of the effect of deployment on the gut microbiome and traveler’s diarrhea susceptibility

**DOI:** 10.1101/2020.07.29.226712

**Authors:** Blake W. Stamps, Wanda J. Lyon, Adam P. Irvin, Nancy Kelley-Loughnane, Michael S. Goodson

## Abstract

Traveler’s diarrhea (TD) is a recurrent and significant issue for many travelers including the military. While many known enteric pathogens exist that are causative agents of diarrhea, our gut microbiome may also play a role in travelers’ diarrhea susceptibility. To this end we conducted a pilot study of the microbiome of warfighters prior to- and after deployment overseas to identify marker taxa relevant to traveler’s diarrhea. This initial study utilized full-length 16S rRNA gene sequencing to provide additional taxonomic resolution towards identifying predictive taxa.16S rRNA analyses of pre- and post-deployment fecal samples identified multiple marker taxa as significantly differentially abundant in subjects that reported diarrhea, including *Weissella, Butyrivibrio*, *Corynebacterium*, uncultivated Erysipelotrichaceae, *Jeotgallibaca*, unclassified Ktedonobacteriaceae, *Leptolinea*, and uncultivated Ruminiococcaceae. The ability to identify TD risk prior to travel will inform prevention and mitigation strategies to influence diarrhea susceptibility while traveling.

## 1 Introduction

Traveler’s diarrhea (TD) is a recurrent and serious issue for many travelers abroad including the military. While not as severe as acute injuries, TD is the most prevalent illness during deployment, and military populations have experienced TD while on deployment since records began (Connor and Farthing, 1999). Deployed populations experience up to 36.3 cases of TD per 100 person-months, with the highest incidence rate during the first month of travel (Olson et al., 2019). Traveler’s diarrhea is commonly ascribed to pathogenic bacteria such as *Salmonella*, *Campylobacter jejuni*, and *Escherichia coli* (including enterotoxigenic, enteroaggregative, and enteropathogenic strains), however, 40 percent of reported cases of TD have no known etiology (Olson et al., 2019). One possible explanation for the unknown or unclassified etiology of TD is a disruption of the collective gut microbiota, or gut microbiome.

The human gut microbiome encompasses thousands of bacterial, eukaryotic, and archaeal species (Huttenhower et al., 2012). A number of human ailments includes an altered or aberrant microbiome. Daily sampling and community analysis revealed how profoundly our gut can be altered by infection as well as diet change during travel (David et al., 2014); however, this study relied on a small number of participants. Military populations are vulnerable to a multitude of stressors including sleep disruption, psychological stress, circadian disruption, significant alterations to diet, and environmental stressors, all of which likely contribute to alterations in the gut microbiota (Karl et al., 2018).

While identification of specific pathogens is certainly of interest, within military populations or any traveler, understanding if a specific microorganism or microbiome population can predict susceptibility or resilience to diarrhea is potentially more desirable to prevent the problem before it could become an impediment to performance. Previously a study within the US Air Force (USAF) at an air base in Honduras identified microbial taxa that were potentially predictive of the development of TD within the study population (Walters et. al 2020). However, in this study samples were collected once the population was deployed and all subjects were deployed to a singular location.

To expand upon previous deployment associated TD work, we initiated a pilot study of the gut microbiome of warfighters both prior to, and after deployment at a multitude of locations across the globe. This pilot study offers an view into the effect of deployment and TD on the microbiome of warfighters through the use of surveys and a sheared 16S rRNA gene sequencing approach to identify potentially predictive microbial biomarkers.

## 2 Materials and Methods

### 2.1 Sampling and Survey of Deployment Associated Diarrhea

Potential subjects were consented and briefed about the study during a recruitment period prior to their individual deployments. After deployment, subjects were also asked to answer a short survey (Supplemental file S1) related to dietary choices, diarrhea before, during, and after deployment, and the number of episodes of diarrhea during each of these three phases. Data were gathered, de-identified, and then recorded digitally with no record of subject name or other identifiable information. Subjects that consented to fecal sampling utilized the BBL CultureSwab EZ II Collection and Transport System (Becton, Dickenson and Company, Sparks, MD) to directly collect a small amount of fecal material after a bowel movement commensurate with the protocol employed by the American Gut Project (McDonald et al., 2018). Collection tubes containing samples were returned within 24 hours of the bowel movement and were frozen at −80 °C. This study was approved by the Institutional Review Board at the 711th Human Performance Wing, Air Force Research Laboratory at Wright Patterson Air Force Base in Ohio, protocol number FWR20150052.

### 2.2 DNA Extraction and Sequencing Library Preparation

DNA was extracted from swabs using the QIAamp DNA Microbiome kit (QIAGEN, Germantown, MD) according to manufacturer’s instructions. The 16S V2-V8 rRNA segments were amplified using PCR with primers S-D Bact 0008 c-S-20 (5’ AGRGTTYGATYMTGGCTCAG3’) and S-D Bact 1391-a-A-17(5’GACGGGCGGTGWGTRCA’3) (Klindworth et al 2013). Briefly, a 50 μL reaction was set up using AccuStart II PCR Supermix (Quantabio, Beverly, MA) per the manufacturer’s instructions, including 10 ng of template DNA. Polymerase chain reaction (e.g. amplicon) cycling parameters were as follows: 3 minutes at 95 °C, then 25 cycles of 30 seconds at 95 °C, 30 seconds at 55 °C, and 30 seconds at 72 °C, with a final extension at 72 °C for 5 minutes. Amplicons were visualized by Invitrogen E-gel agarose gel electrophoresis, and then purified using Agencourt AMPure XP beads at a final concentration of 1.8 X (Beckman Coulter, Brea, CA). Final amplicon concentration was quantified using Qubit dsDNA high sensitivity assay (Life Technologies).

Amplicon sequencing libraries were then prepared using an Illumina Nextera XT Library preparation kit (Illumina, San Diego, CA), followed by sequencing PE150 sequencing on a Illumina MiSeq.

### 2.3 Data analysis

Sequence reads were assigned initial taxonomy using Kraken 2 (Wood and Salzberg, 2014) with the SILVA database (Quast et al., 2013) serving as a reference. After initial assignment taxonomy was further refined using Bracken (Lu et al., 2017) for each sample. Individual Bracken reports were then imported into Pavian (Breitwieser and Salzberg, 2016) before a taxonomy table containing all samples was exported as a tab delimited file. This was then imported into Phyloseq within R (McMurdie and Holmes, 2013; R Core Team, 2013) to aid visualization and statistical testing. Ordinations were generated within AmpVis2 (Andersen et al., 2018). A Dirichlet Multinomial Mixture Model was also produced to estimate the number of metacommunities within the dataset (Holmes et al., 2012) using the microbiome R package. Differential abundance and variability was assessed using COunt RegressioN for Correlated Observations with the Beta-binomial, or Corncob (Martin et al., 2019)

## 3 Results

A total of 521 subjects responded to survey questions, and 23 provided fecal samples either prior to or after deployment, or both. A total of 56 fecal samples were collected from the cohort of 23 subjects with 37 collected prior to deployment and 19 after return from deployment. Subjects reporting diarrhea on deployment ranged from 10 percent in North America (NORTHCOM) to 67 percent in CENTCOM (Figure 1a). Of those reporting diarrhea, multiple instances were common, again mostly within CENTCOM (Figure 1b). Additional survey results are available in supplemental data (Supplemental Table S1). A total of 52 percent of subjects that donated feces reported diarrhea on deployment, in line with survey statistics.

**Figure 1:**
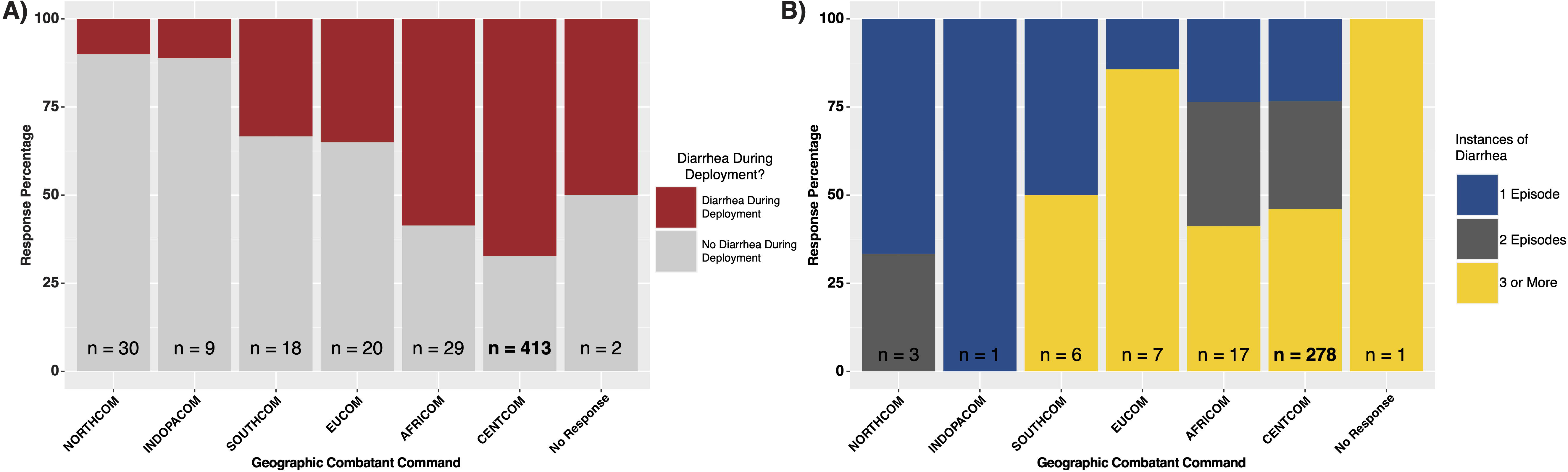
Deployment diarrhea survey summary statistics. (A) ‘yes/no’ response grouped by geographic combatant command, and (B) for those that responded ‘yes’, the number of instances of diarrhea on deployment, again grouped by combatant command.

All 56 samples were successfully sequenced resulting in 5.5 million sequence reads. Median library size was 103,994 reads after quality control and taxonomy assignment. Individual library size and metadata can be found in supplemental table S2. Overall there was a non-significant difference between pre- and post-deployment samples (ADONIS p > 0.05) although individual subjects with linked pre- and post-deployment samples, particularly Subjects 2 and 34, had large shifts in community membership (Figure 2). After assignment of samples into metacommunities by DMM, general trends of community membership emerged (Figure 3a). These included a high Bacteroidetes relative abundance group (Metacommunity 1), an intermediate Bacteroides group (Metacommunity 2), and a low Bacteroides/high Prevotella group (Metacommunity 3). Subjects 2 and 34 had an opposite shift in the relative abundance of the *Escherichia/Shigella* present (Figure 2) before or after deployment, with a notable decline from 12.8 to 0 percent for subject 34, and an increase from 0.1 to 22.6 percent relative abundance for subject 2. Neither subject experienced diarrhea on deployment. Many subjects remained within the same metacommunity before and after deployment, although several (S2, S4, S5, S6) shifted between a high and intermediate *Bacteroides* relative abundance (metacommunities 1 and 2) (Figure 3a). Overall diversity of a subject’s gut microbiota was also very similar before and after deployment (Figure 3b), and was not significantly different (p > 0.05). This is supported by a relatively low number of taxa (17, Fig. 4b) having differential abundance before or after deployment. However, if samples are tested for differential abundance of taxa by pre/post deployment status and those that reported diarrhea on deployment, 32 taxa were differentially abundant towards those with diarrhea (Figure 4a). The majority of these taxa were very low relative abundance (Supplemental Figure S1a and b); however one taxon, Ruminococcaceae UCG-014, was both differentially abundant in those with diarrhea during deployment and above 3 percent relative abundance on average in all sampled communities prior to deployment (Figure 3a). Other strongly differential taxa included *Weissella, Butryvibrio, Brenneria, Buchnera,* and *Sutterella* (Figure 3a). Taxa differentially abundant before or after deployment included *Lactobacillus, Pantoea, Cornybacterium, Megamonas, Brenneria,* and *Anaeroplasma* (Figre 4b). While significantly differentially abundant taxa were observable between sample grouping, both by diarrhea status (yes vs. no) on deployment and by deployment status (pre- vs. post deployment), abundances of most differential taxa were largely invariant within sample groups (Supplemental Figure S2).

**Figure 2:**
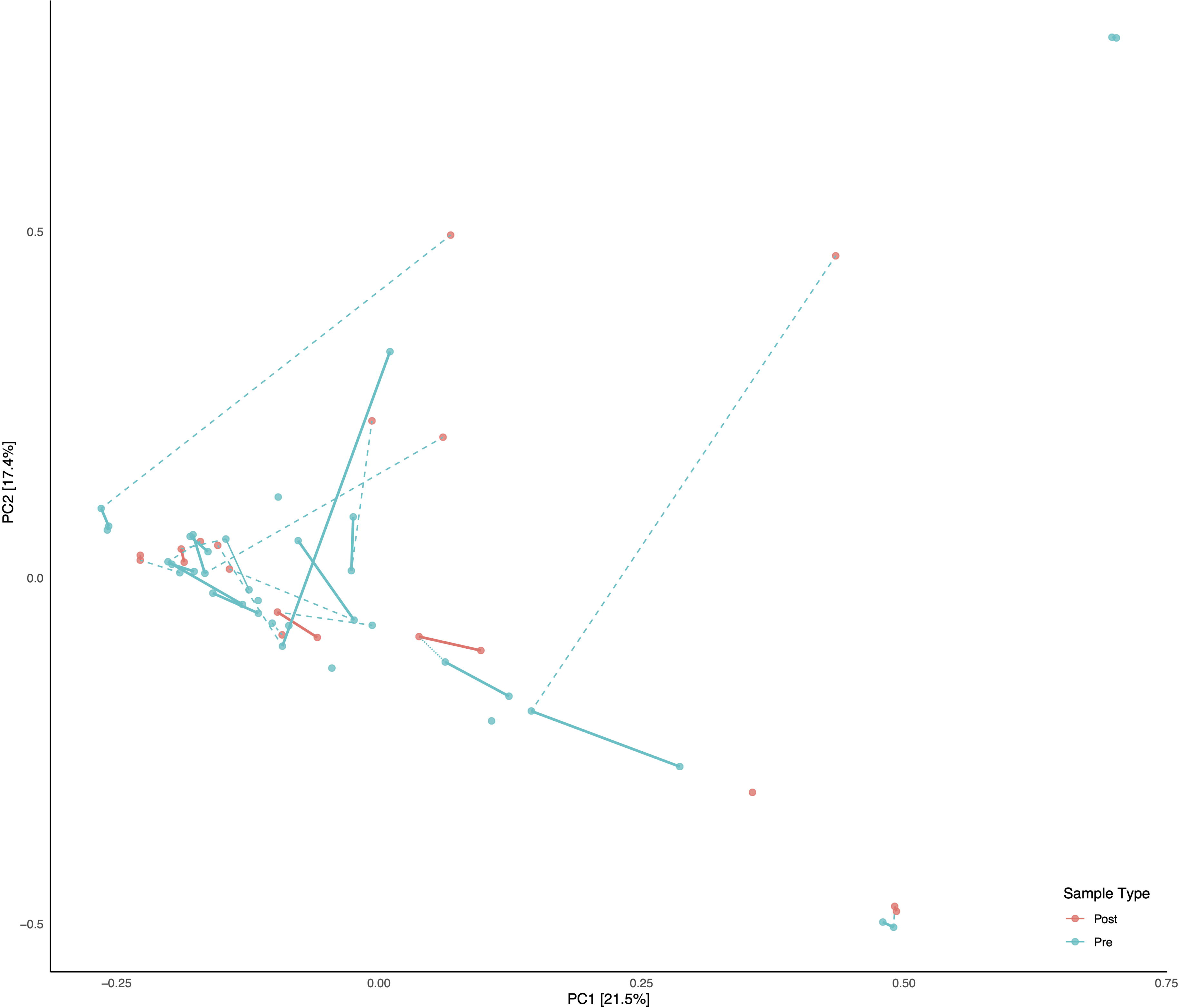
Principal Component Analysis (PCA) of all samples. Subjects with linked pre/post samples are shown with dashed lines connecting the pre and post sample types. Solid bars connect the same sample types (Pre or Post) within each subject on the PCA.

**Figure 3:**
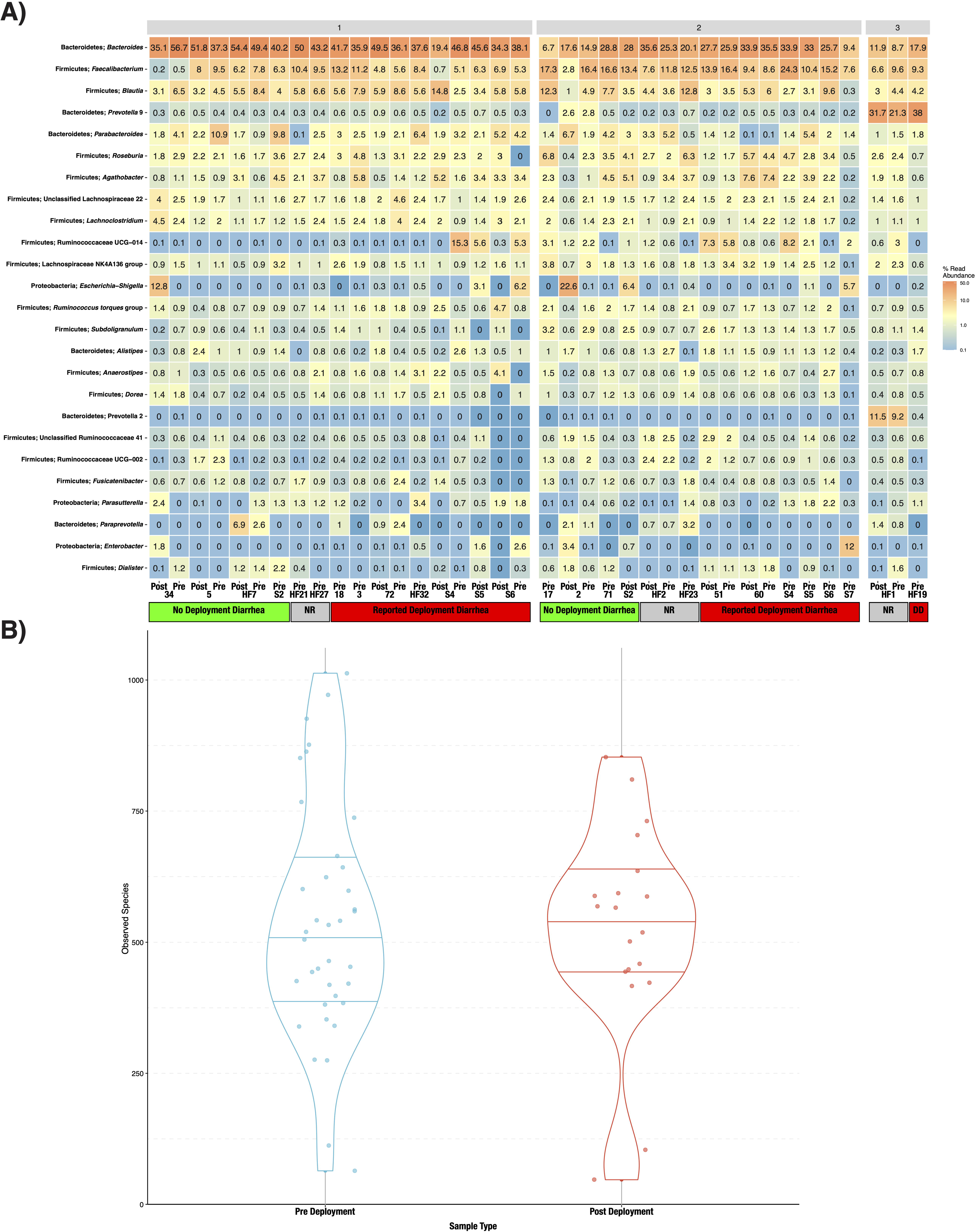
Heatmap of the 25 most abundant taxa, clustered by genus (A). Samples are faceted by DMM metacommunity, and by whether the respondents were diarrheal prior to, and while on deployment. Subjects that gave no response to the questionnaire are noted as “unknown” within the heatmap. (B) Observed alpha diversity, estimated by number of species in samples before and after deployment.

**Figure 4:**
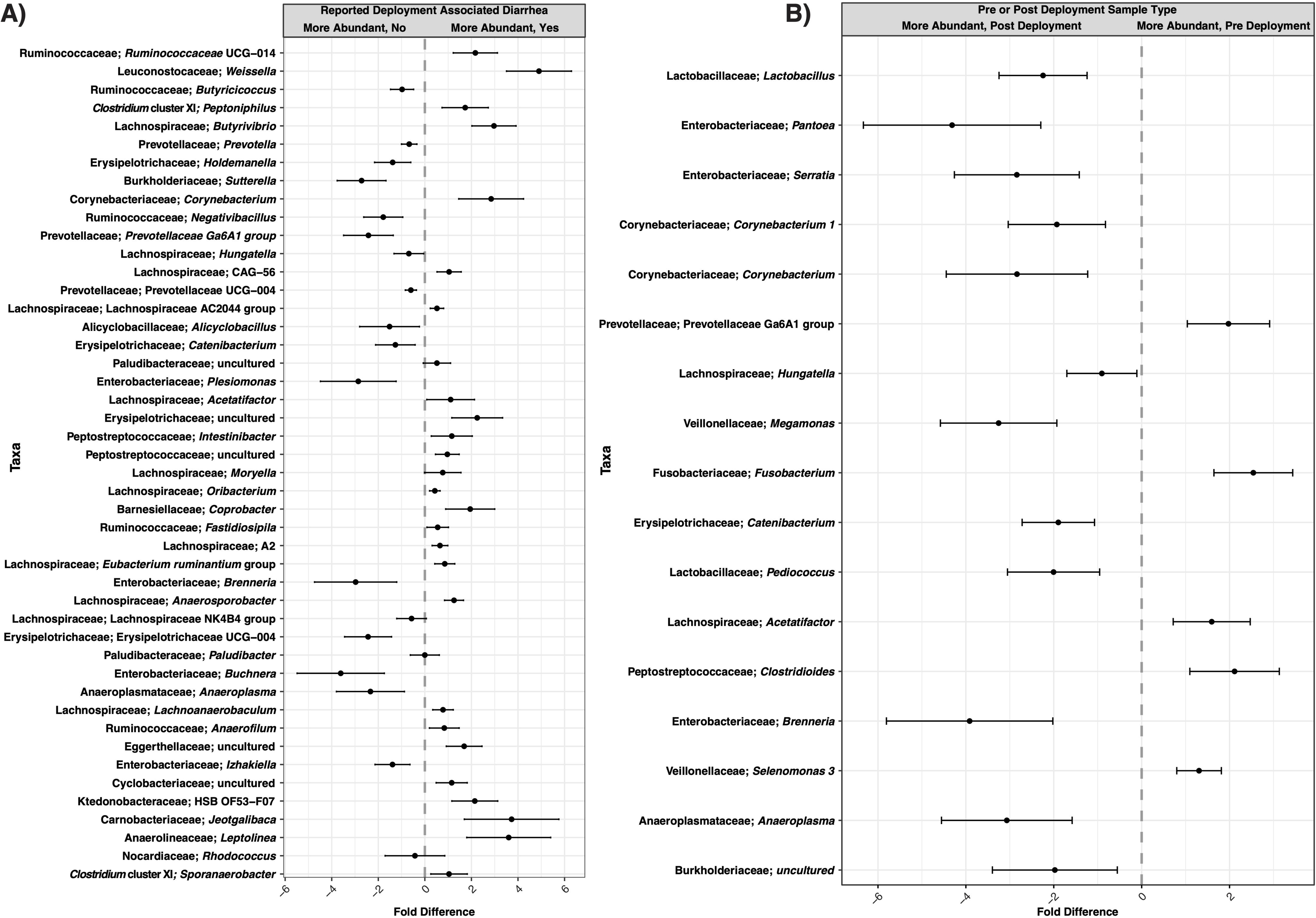
Differentially abundant taxa in subjects with linked (pre/post deployment) samples, either comparing subjects with or without diarrhea on deployment (A) or those taxa differentially abundant before and after deployment (B). Only taxa found to be significantly differentially abundant after false discovery rate error correction are shown in the figure.

## 4 Discussion

TD impacts over half of individuals during deployment, representing a detriment to personal comfort, health, and the ability to carry out mission objectives. A recent comprehensive review of TD literature by Olson *et. al* (Olson et al., 2019) suggests that acute TD is still prevalent among not just the military but the broader American population as well. Any ability to predict if certain individuals may be more susceptible than others to TD would greatly improve the performance of military travelers and also improve the quality of life of all individuals traveling abroad by ensuring individuals take more care in food choices, administration of prophylactic TD medicines, and more broadly by being more aware of their individual risks for diarrhea while abroad. This work identified several potential marker taxa worthy of additional study in the future, with more comprehensive deep metagenomic sequencing studies. Further, we have shown the need for sampling not just after diarrhea or other negative effects have occurred, but in collection of samples prior to deployment.

With few exceptions, prior to, and after, deployment the subjects were separated into three distinct metacommunities identified by utilizing the Dirchelet Multinomial Mixture modeling (Figure 2). Largely these communities were influenced by two major gut bacterial genera, the *Bacteroides* and the *Prevotella*. Notably, *Prevotella* were largely absent within the two other major communities (Figure 3). Previous large-scale analyses have noted that *Prevotella* exist as one end-member (with *Bacteroides* as the other) of a gradient of human gut microbiomes, rather than discrete enterotypes (Jeffery et al., 2012). Our work appears to support this assertion. Overall the literature is mixed in the potentially dys- or eubiotic properties of *Prevotella* (Precup and Vodnar, 2019). However, increased relative abundances of *Prevotella* are associated with increased risk of diarrheal irritable bowel syndrome, or IBS-D, while individuals with high relative abundances of *Bacteroides* were not (Su et al., 2018). Within this study only two subjects were within the *Prevotella* dominated community, and only one completed a survey, but did report diarrhea while on deployment.

Focusing instead on subjects whom reported diarrhea status while on deployment and also donated linked samples before and after deployment, we can identify several microbial taxa of interest for further study. Differential abundance analyses were used to determine which, if any, microbiota may be linked causally to predict if a subject were to develop or be protected from diarrhea during deployment. Limiting the focus to those taxa identified as being more than two fold differentially abundant, seven taxa were associated with a lack of diarrhea, including the Enterobacteriaceae *Buchnera, Brenneria,* and *Plesiomonas*; as well as *Sutterella,* Prevotellaceae Ga6A1, Erysiplotrichaceae UCG-004, and *Anaeroplasma*. Both *Buchnera* and *Brenneria* are more associated with plants, either as obligate endosymbionts or as pathogens with little to no known association with the human gut (Douglas, 1998; Octavia and Lan, 2014). *Plesiomonas* was previously associated as a potential, emerging enteric pathogen (Theodoropoulos et al., 2001) and so its’ association within our dataset with subjects that did not develop diarrhea requires further study. Members of the Prevotellaceae, specifically the *Prevotella* are associated with vegetarian or vegetable rich diet (Precup and Vodnar, 2019). The specific Prevotellaceae we identified, Ga6A1, was previously associated with cellulose rich diets in waterfoul (Guo et al., 2019). The Erysiplotrichaceae have conflicting associations, and while increased relative abundances of this family have been associated with gut inflammation, the opposite has also been found (Kaakoush, 2015).

Taxa that were increased in subjects reporting TD included *Weissella, Butyrivibrio, Corynebacterium,* uncultivated Erysipelotrichaceae, *Jeotgallibaca,* unclassified Ktedonobacteriaceae and *Leptolinea*. *Weissella* and *Butyrivibrio* were previously reported as either beneficial (via production of butyrate) or as potential probiotics so their increased abundance is curious, although literature related to *Weissella* is unsettled on its potential pathogenicity, and of interest in further study (Abriouel et al., 2015; Vital et al., 2017). The Erysipelotrichaceae are highly immunogenic and may contribute to gut inflammation and/or gastrointestinal issues (Kaakoush, 2015). Both the Ktedonobacteriaceae and *Leptolinea* are within the Chloroflexi, a highly diverse lineage adapted to a multitude of diverse aquatic and terrestrial environments including the mammalian gut, although at a low relative abundance (Ley et al., 2008). While not more than two-fold differentially abundant, members of the Bacteroidetes included an increase in abundance of the genus *Coprobacter* in subjects that reported diarrhea during deployment. *Coprobacter* was previously associated with a Chinese population that consumed a largely high fat diet (Qian et al., 2018) but, to date, we could find no prior association of *Coprobacter* with any specific disease status.

While most of these differentially abundant taxa were not high in relative abundance (Figure 3, supplemental table S3), one unclassified genus within the *Ruminococcaceae* family was identified. *Ruminococcaceae* UCG-014 relative abundance was differentially abundant in subjects that reported TD during deployment (Figure 4) and was in higher relative abundance as well (Figure 3). *Ruminococcaceae* UCG-014 has been previously described in mouse model studies of neuropathic pain and in cattle microbiome surveys, but to this point never associated as a potential predictive marker of diarrhea (Yang et al., 2019; Zhang et al., 2019). Within metacommunity 1, UCG-014 was more abundant prior to deployment; however, in metacommunity 2 this pattern was flipped and a greater relative abundance of the genera was found in post deployment samples. Still, it could be that the lineage occupies a fundamental niche in the guts of those predisposed to TD. Other unclassified Ruminococcaceae genera have been recently associated with traveler’s diarrhea, specifically Ruminococcaceae UCG-013 which was differentially abundant in a deployed military population in Honduras (Walters et. al 2020). Further work will confirm the role unclassified Ruminoccocaceae play in the development of diarrhea, but their presence could predispose those in metacommunity 2 to TD in future deployments. There is certainly anecdotal evidence that gastrointestinal issues follow multiple TD episodes during deployment, but this correlation requires additional study. Future, larger investigations involving deployed populations will attempt to further elucidate the connection between *Ruminococcaceae* UCG-014 and the risk of diarrhea, if any.

The identification of microbial risk factors associated with TD remains of crucial interest to military populations worldwide. Our work represents a first step towards larger, ongoing investigations of deployed military populations. Caution is warranted in distilling these results to provide actionable information. The number of subjects that completed both pre and post sampling was small (n = 13) limiting the power of the study to produce definitive conclusions. Future studies will include larger cohorts and follow their performance and diet prior to, during, and after deployment to more conclusively identify microbiota correlated to diarrheal status, with particular interest towards the Ruminococcaceae. This preliminary study suggests that several taxa may be predictive of diarrhea status prior to travel. The ability to identify TD risk prior to travel has benefits beyond the military and will inform prevention and mitigation strategies for the comfort of any traveler in the future, including modulation of the gut microbiome using prebiotic, probiotic, or synbiotic methods.

## 5 Conflict of Interest

The authors declare that the research was conducted in the absence of any commercial or financial relationships that could be construed as a potential conflict of interest. BWS is an employee of UES, Inc.

## 6 Author Contributions

BWS conducted the analyses and prepared the manuscript. MSG conceptualized the experiment, conducted sampling, and edited the manuscript. WL generated the sequencing libraries and edited the manuscript. AI conduced sampling and edited the manuscript. NKL conceptualized the experiment and edited the manuscript.

## 7 Funding

This work was made possible by the 711th Human Performance Wing Research, Studies, Analyses, and Assessment Committee Intramural Grant

## 8 Acknowledgments

We would like to acknowledge the Airman and Family Readiness Center, Wright-Patterson AFB as well as the Airman Systems Directorate Technical and Staff Sergeants whom assisted in numerous ways throughout the study. This manuscript was approved for public release on 28 July 2020, Distribution A, distribution unlimited; PA case number 88ABW-2020-2343; MSC/PA-2020-0173.

## 9 Data Availability Statement

The dataset generated for this study can be found in the NCBI SRA at XXXX.

## 11 Figure Legends

Supplemental Figure S1 Heatmap of significantly differentially abundant taxa (A) in subjects with traveler’s diarrhea during deployment and significantly differentially abundant taxa in subjects before and after deployment (B).

Supplemental Figure S2: Differential variability of taxa in subjects with linked (pre/post deployment) samples, either comparing subjects with or without diarrhea on deployment (A) or those taxa differentially abundant before and after deployment (B). Taxa shown are only those found to be differentially abundant in figure 4.

Supplemental Table S1: Detailed survey responses

Supplemental Table S2: Mapping file including read numbers per library

Supplemental Table S3: Per sample taxonomy table after merger in Pavian

