## Supplementary figures and images for "A pilot study of the effect of deployment on the gut microbiome and traveler’s diarrhea susceptibility"

### Figure S1

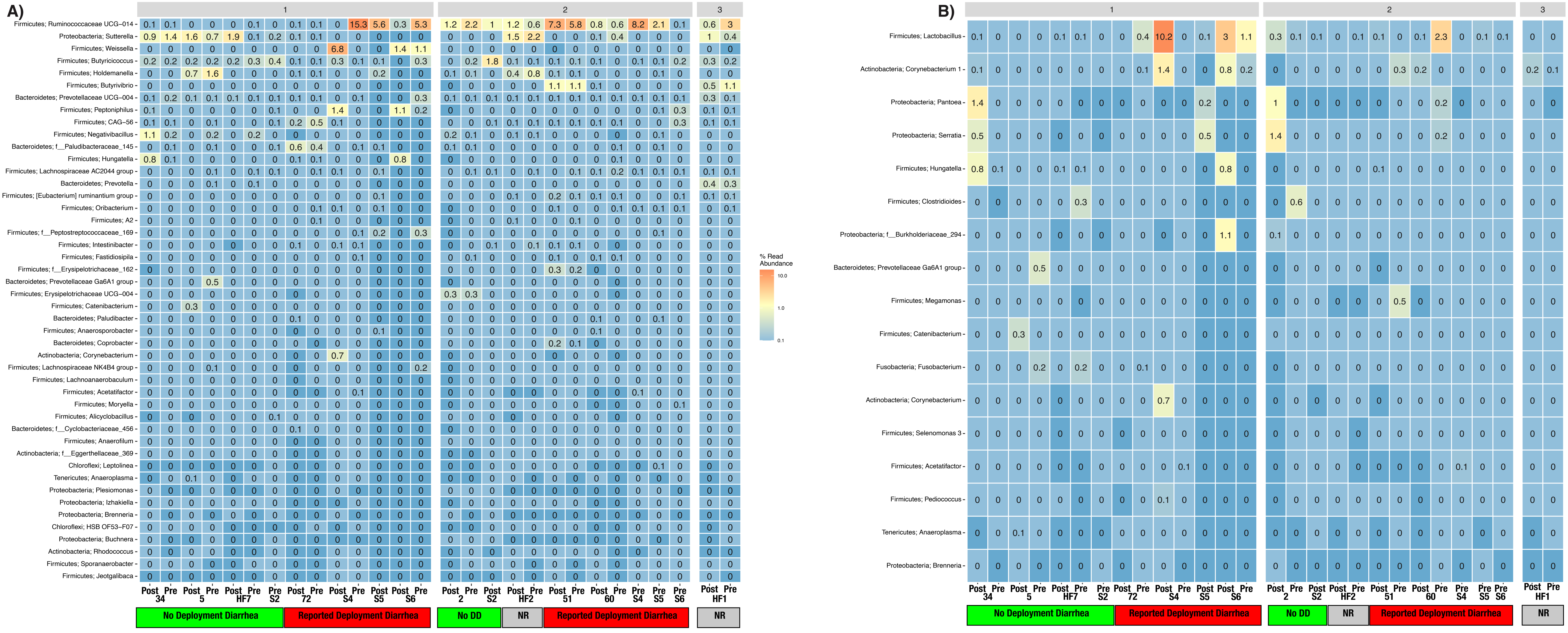

### Figure S2

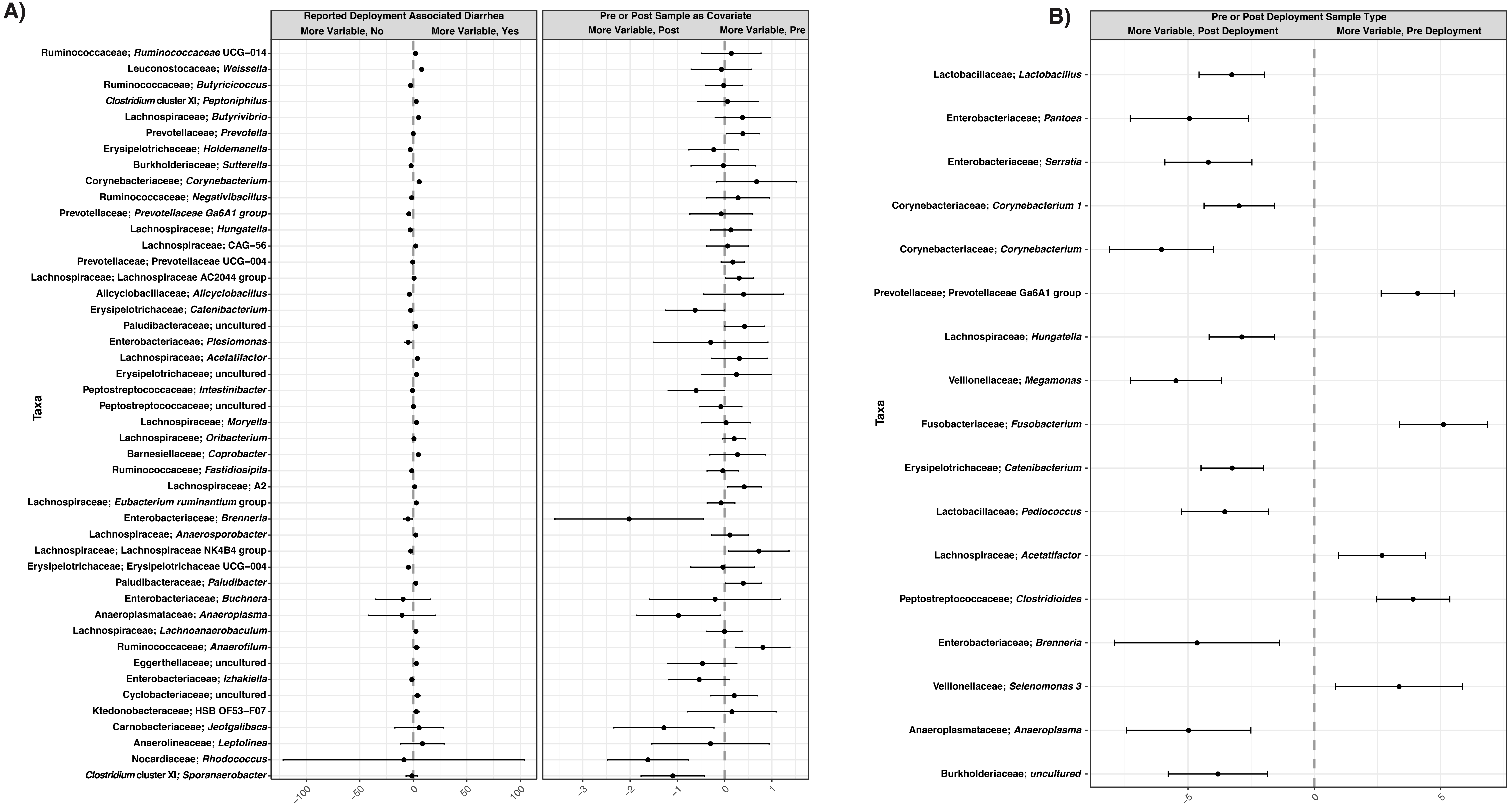
